# Scaffolded Learning of Bipedal Walkers: Bootstrapping Ontogenetic Development

**DOI:** 10.1101/2020.10.03.324632

**Authors:** Jiahui Zhu, Chunyan Rong, Fumiya Iida, Andre Rosendo

**Affiliations:** Living Machines Lab, School of Information Science and Technology, ShanghaiTech University; Bio-Inspired Robotics Lab, Department of Engineering, University of Cambridge

**Keywords:** Bipedalism, Ontogeny, Scaffolded Learning, Bootstrapping

## Abstract

Bipedal locomotion has several key challenges, such as balancing, foot placement, and gait optimization. We reach optimality from a very early age by using natural supports, such as our parent’s hands, chairs, and training wheels, and bootstrap a new knowledge from the recently acquired one. In this paper, we propose a scaffolded learning method from an evolutionary robotics perspective, where a biped creature achieves stable and independent bipedal walking while exploiting the natural scaffold of its changing morphology to create a third limb. Hence, we compare three conditions of scaffolded learning to reach bipedalism, and we prove that a performance-based scaffold is the most conducive to accelerate the learning of ontogenetic bipedal walking. Beyond a pedagogical experiment, this work presents a powerful tool to accelerate learning on robots.

## I. INTRODUCTION

Compared to wheeled and multi-legged robots, bipedal robots are more adaptable to complex terrain, have higher mobility, and receive increased attention from researchers. However, bipedal walking [1] is a notoriously tricky task due to its inherent instability and dynamics. In [2] seven intrinsic difficulties are listed, such as non-linear state space, the effect of gravity, the influence of semi-structured and unknown environments, nominally instability, multiple inputs and multiple outputs, dynamic characteristics over time, and requiring continuous and discrete control. In the face of such challenges, the design of physical prototypes and complex control algorithms for bipedal walking is costly and time-consuming, and these will continue to be the main bottleneck for the deployment of affordable bipedal robots [3].

Scaffolding is a learner-centered teaching method based on the constructivist learning theory, aiming at cultivating students’ problem-solving ability and autonomous learning ability. Pedagogy explains it as providing small-step clues or hints (scaffolds) for students to learn step by step to discover and solve problems gradually. This method leads to students mastering the knowledge to be learned, improving their problem-solving ability, and eventually growing into independent learners. Vygotsky, a famous psychologist in the former Soviet Union, derived this teaching idea from the “zone of proximal development” theory [4]. Pedagogical applications have actively used scaffolding to bootstrap knowledge [5] [6], and some researchers tried to understand its associations with human locomotion: the ontogenetic development [7] of bipedal walking in human infants [8], and the mechanism of acquiring general motor skills and of human walking [9] [10]. Besides, reaching bipedal locomotion during early childhood requires individuals to be strong enough to support their weight, stable enough to resist unstable gravitational force, and to move in a state of dynamic balance when the body alternates between double support and single support [11] [12]. Along these lines, the work from [13] simulated a supported infant walking by applying linear springs and dampers to a bipedal walking model, which was controlled by a rhythm-generation mechanism called central pattern generator (CPG) [14]. As this simulated infant learned to walk, it naturally became upright and prescinded from the spring support, but A. the mechanisms deciding the amount of/need for supports to be given or B. the benefits compared to the absence of such supports were not investigated.

In this article, we present a scaffolded learning method for bipedalism to bootstrap its ontogenetic development with gradual morphological and control changes. Inspired by the mechanisms of a child learning to ride a bike with training wheels, as shown in Fig. 1, our simulations consist of a two-legged creature with a long body that can be dragged through the floor as a tripod, providing a scaffold during the learning of bipedalism. In addition, we use a genetic algorithm [15] to evolve both morphology and control (virtual model control [16]). We define the fitness function as the walking distance in fifteen seconds divided by the leg length. In our initial simulations, the creature tries to maximize its walking speed while evolving freely, and later we introduce two evolving conditions where we force the creature to reduce its body length (abdicate from its scaffold) over fixed time intervals or once the creature makes gradual performance improvements. We show that the performance-based scaffold is superior to the time-constrained case and the free scaffolded case, as it allows the controller to mature before becoming independent from the morphological support. We explain the effectiveness of the bootstrapping mechanism, draw parallels to robotic implementations of ontogeny [17], and propose a framework where real-world robots can use a similar approach to bootstrap knowledge [18]. In Section II, we describe our adopted methods and propose the performance metrics for our simulation, along with three scaffolded learning cases. We show the results in Section III and discuss the implication of these results in Section IV. We conclude our work in Section V.

**Fig. 1:**
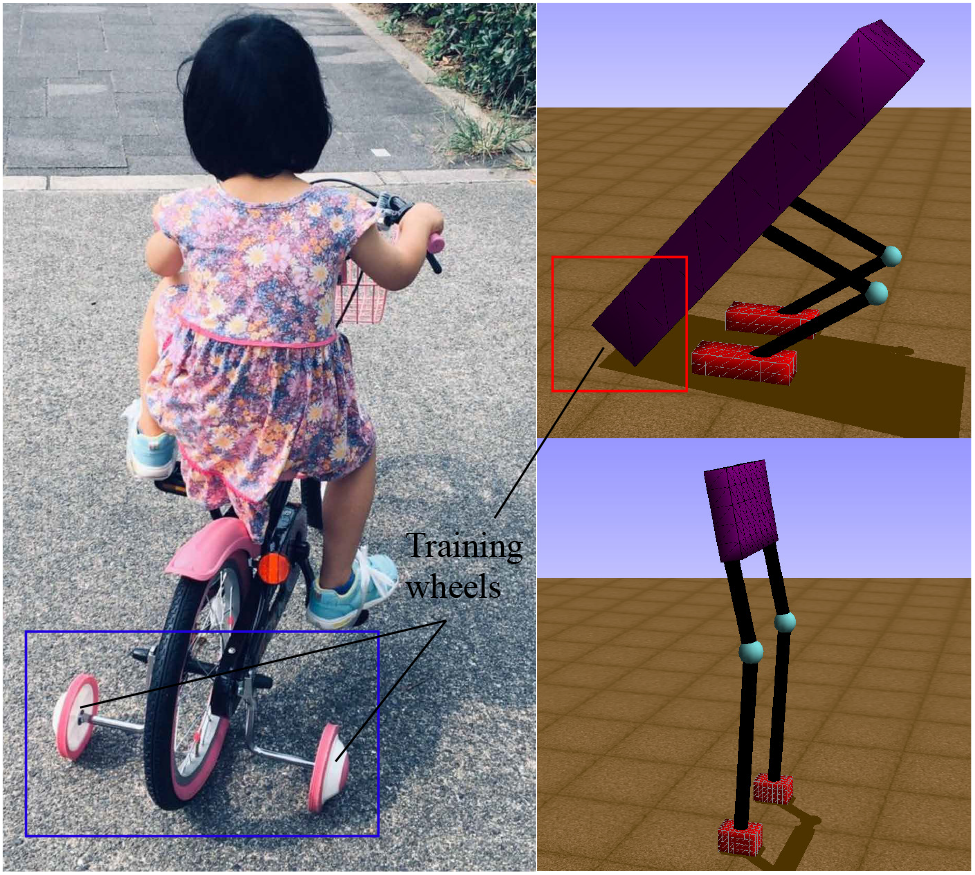
The general idea of scaffolded learning can be exemplified with a child on training wheels. A biped creature’s body length can be adjusted to act as a tripod with shorter legs, sacrificing speed to gain stability, while a longer leg and short body would be optimal bipedalism.

## II. Material and Methods

### A. Genetic Algorithm

Genetic Algorithms (GA) [15] use simulated evolution to search for solutions to complicated problems. The algorithm works applying selection, recombination, and mutation processes on encoded genotypes, and evaluate the fitness of each individual to evolve it over generations. We started GA with a default individual in our work, which has a long body touching the floor as a support, and then we recombined and mutated it to produce a generation. We adopted a subset of the generation containing the fittest individuals to create the new generation. The algorithm used an array of individuals to remember the fittest ones ever produced and utilized these to mutate new generations under an exploration-exploitation trade-off so that GA could find a globally optimal solution. The pseudo-code of this algorithm can be found in Algorithm 1.

### B. Virtual Model Control

Virtual Model Control (VMC), developed by Pratt et al. [2] [16], is a motion control framework that uses a desired virtual force, combined with the kinematics model and its Jacobian, to generate desired joint torques on the stance leg. The combination of these joint torques creates the same output that the virtual force would have created, thereby creating the intended motion on the robot/creature. Such forces can be emulated as products from many components, such as springs, dampers, dashpots, masses, dissipative fields, or any other imaginable component.

Genetic Algorithm

**Algorithm 1.**
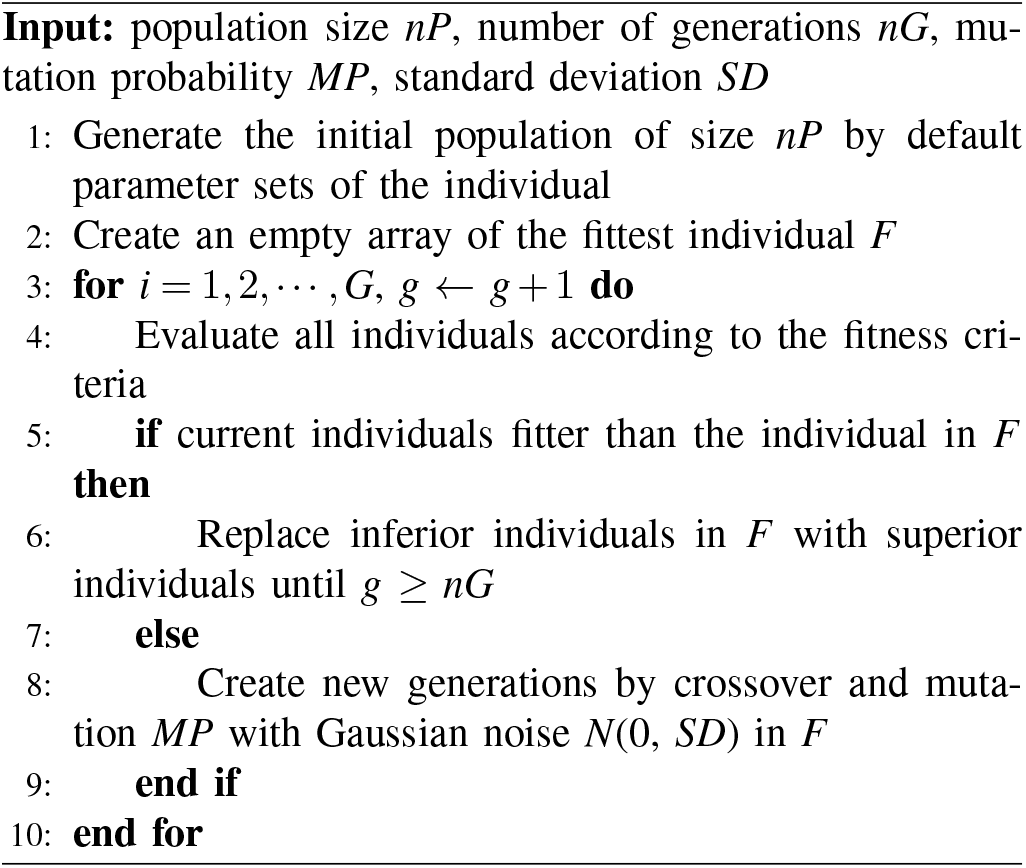

Here, we applied VMC as an intuitive controller algorithm of the tripod creature to perform a forward walking with horizontal, vertical, and torsional springs, and dashpots components, as shown in Fig. 2. VMC’s benefits are that it is compact, requires relatively small amounts of computation, and can be implemented in a distributed manner. We could also implement a high-level controller as a state machine that changes virtual component connections or parameters at the state transitions. Even though we use a discrete high-level controller, the overall motion can be smooth if the virtual components have a low-pass filter effect.

**Fig. 2:**
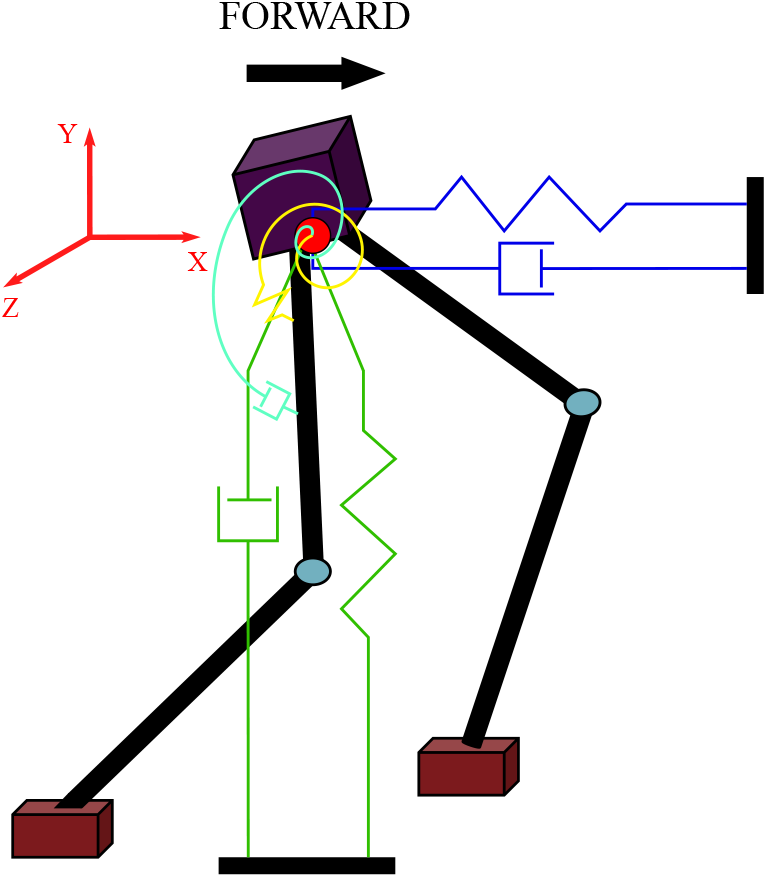
The virtual model in the simulation. We attach linear springs and dashpots to the individual’s hip position as the granny walker mechanism, proposed by [2], to maintain a constant height, and the dog-track bunny mechanism applies a virtual force in the forward horizontal direction to help obtain the desired velocity. Also, it has a torsional spring and a rotary dashpot acting on the hip joint to keep the upper body straight in the standing phase, while in the step phase, the hip joint only has a torsional spring to swing the leg.

### C. Walking State Machine

We chose a finite state machine to allow transitions between different virtual models during the walking cycle. It allowed the controller to use a virtual model suitable for the individual’s current position and the walking cycle phase.

Fig. 3 shows the finite state machine used in the algorithm, and the Table I shows the transitions of three states. The first state is the standing phase. When the individual is stable, a test is done to see which foot is in front of the other. If the left foot is in front, then we move into the left support state. In this state, the step phase virtual model is activated. When the right foot moved in front of the left foot and was close to the ground, the state machine progressed to the step phase of the right support state, in which torques in the swing leg became zero. Then, when the right foot landed on the floor in front of the left foot, the state machine returned to the double support state’s standing phase.

**Fig. 3:**
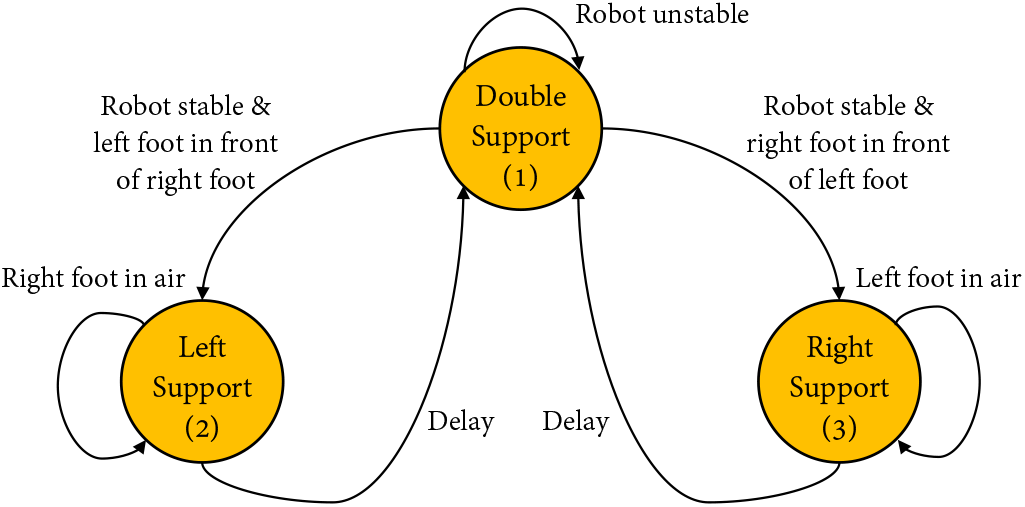
The walking state machine [19] in the genetic algorithm. There are three states during the walking cycle to allow transitions of different virtual models. The first state is double support, which means both feet are contacting the floor, and the second state of left support means only the left foot contacting the floor. Similarly, the third state right support only has the right foot on the floor.

### D. Simbody Simulation Environment

Simbody is a high-performance, open-source C++ library providing sophisticated treatment of articulated multibody systems with particular attention to biomedical simulations’ needs. It is useful for predictive dynamic simulations of diverse biological systems such as neuromuscular biomechanical models and coarse-grained biomolecular modelling. It is also well suited to related simulation domains such as robotics, avatar simulations, and controls, and provides realtime capabilities that make it useful for interactive scientific simulations and virtual worlds [20].

**TABLE I:**
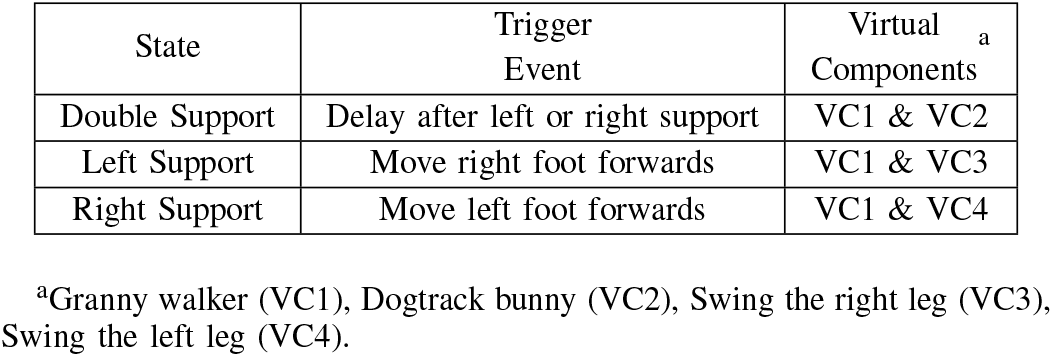
Transitions of Walking State Machine

The simulation was conducted in a DELL OptiPlex 7060 series desktop with Ubuntu 18.04 system, i7-8700 processor, 12 threads, and 32GB internal memory. We will terminate the learning process of bipedal walking early if the individual fell, while the upper body fell to below one half of the total body height, and they overran a time limit of fifteen seconds.

### E. Performance Metric

Our ultimate goal in this work is to evolve independent bipedal walking in fifteen seconds duration simulation, where the fitness is equal to the walking distance divided by the total leg length. Here, we referred the walking Froude number [21], which is the ratio of the centripetal force and the weight of the animal walking:

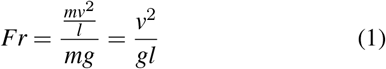

where *m* is the mass, *l* is the characteristic length, *g* is the acceleration due to gravity and *v* is the velocity. The characteristic length *l* is chosen to be the total leg length.

Since we start the simulation at a supported tripod walking, there will be tripod walking individuals, bipedal walking individuals, and even individuals with alternating gaits during the ontogenetic development. We use this behaviour as a metric to measure the performance of bipedal walkers. Also, we observe the growth rate of the fitness and the degree of body length decay as the other two performance metrics. We will conduct the following three cases in different body length constraints in 4000 generations. To verify our results’ repeatability and reliability, we decide to do three replicates for each case.

1. *Free body length scaffold:* According to the current fittest, the genetic algorithm will choose the appropriate combination of body parameters and control parameters, benefiting from the algorithm’s advantages. Therefore, we let it evolve freely without any restrictions.
2. *Time-constrained body length scaffold:* To speed up the evolution of independent bipedal walking, we have artificially restricted the body length by shortening the maximum range of body length proportionally as the generations increase.
3. *Performance-based body length scaffold:* Considering the limitation of the number of generations, it is likely that bipedal walkers’ learning cannot be better explored. Therefore, we have balanced exploitation and exploration and designed a brace that limits the body length according to the current performance, which allows it to explore better at the beginning and focus on exploitation when achieving better results to maximize performance.

## III. Results

We started with a comparison between the best biped and tripod’s fitness from our simulations, as shown in Fig. 4(a). Although these two presented a very similar fitness initially, the biped gradually outperformed the tripod from the four-second mark on-wards and reached a fitness value 37.5% larger than the tripod. We then generated their body trajectories and plotted snapshots of these two creatures walking for five seconds (instead of the total fifteen seconds), as shown in Fig. 4(b). From the figure, the stride length from the best biped was gradually increasing while the best tripod’s stride length was almost constant. In Fig. 5, we demonstrated a variety of tripodal and bipedal creatures obtained during generations, and all their body lengths were gradually decreasing. Interestingly, we observed that most tripod creatures had long and broad feet, while the feet of bipedal creatures were continuously shrinking over generations.

**Fig. 4:**
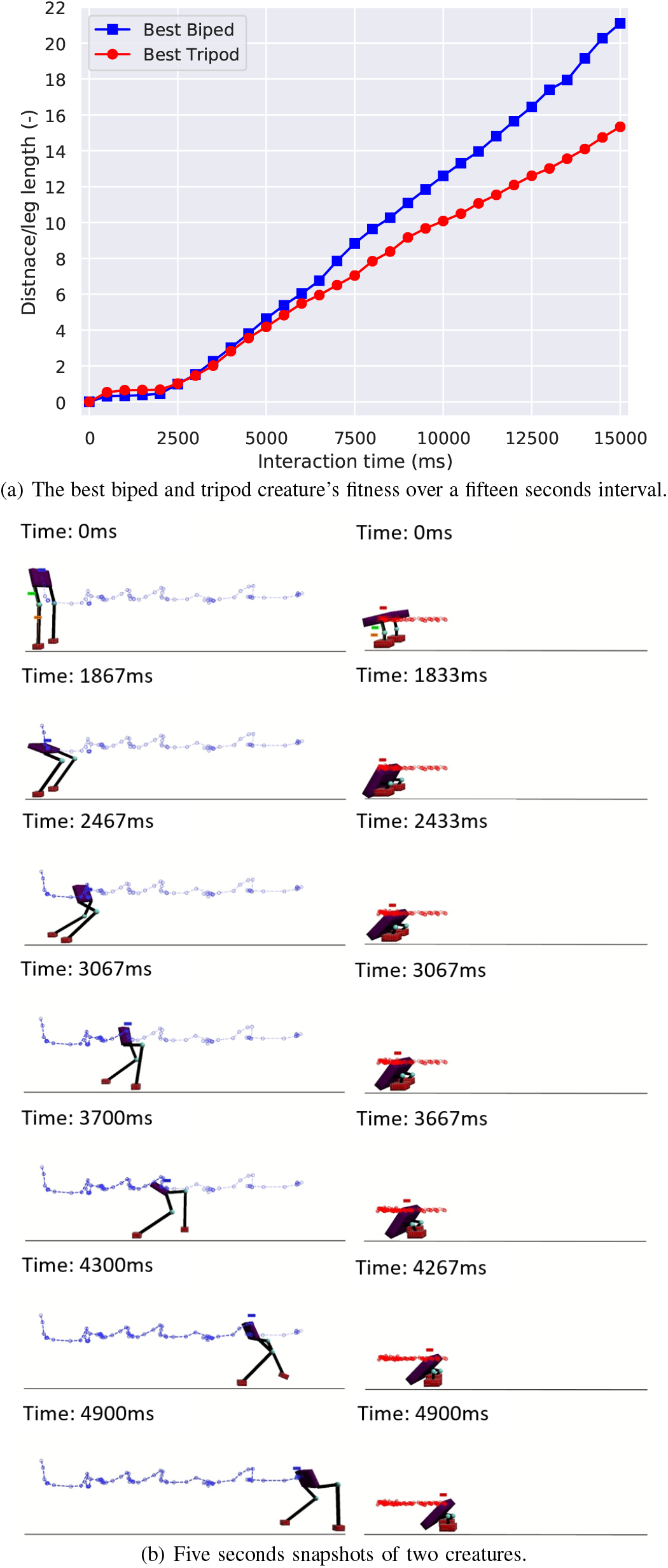
(a) Fitness of the best biped and best tripod creatures. The best biped has a femur length of 0.36 *m* and a tibia length of 0.81 *m*, while the values for the best tripod are 0.20 *m* and 0.21 *m*. (b) The trajectory is generated by the motion analysis, with blue and red lines showing the path of the center of the body.

**Fig. 5:**
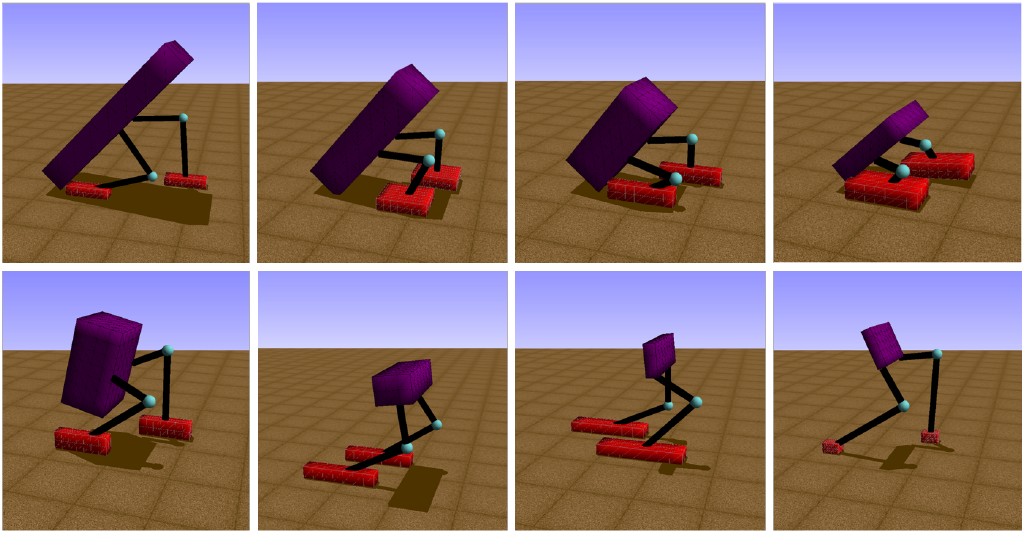
Individuals of tripods and bipeds with different leg lengths, body sizes and foot sizes evolved in the simulation. The upper figure of individuals performed supported tripod walking gaits, and the lower figure of individuals are all bipedalism.

We obtained the gait information from the relative Center of Gravity (CoG) position, the relative horizontal velocity, and the relative vertical velocity for both creatures, shown in Fig. 6. Here, we defined the relative displacement unit as a leg length (*ll*), and the unit of the relative velocity as a leg length per second (ll/s) to fairly compare the locomotion from the best biped and the best tripod. Therefore, as for the CoG, the best biped first fell to −0.35 *ll* and then returned to 0 *ll*, finally oscillating at the original position. The best tripod first fell to −0.2 *ll* and oscillated for the first eight seconds. Then, it fell again to −0.3 *ll*, gradually rose back to −0.15 *ll*, and stabilized at that height. About the relative horizontal velocity, the best biped creature reached a maximum of 8 *ll/s*, while the best tripod creature merely obtained 6 *ll/s*, so the relative horizontal velocity from bipeds was 30% higher than tripods. Regarding the relative vertical velocity, the best biped creature reached a maximum of 2 *ll/s* while the best tripod creature was half of it, at 1 *ll/s*. We plotted the variations that happened with body, femur and tibia length throughout the evolution of the bipedal creature, and we show it at Fig. 7. The body length gradually decreases from 0.9 *m* to 0.1 *m* over 1000 generations, while the femur and tibia lengths had very few changes in this interim. After 1000 generations, the body length continued decreasing, reaching 0.05 *m* (the minimum value set in the simulation), while the tibia length increased 0.15 *m* and the femur length kept oscillating stably at 0.4 m.

**Fig. 6:**
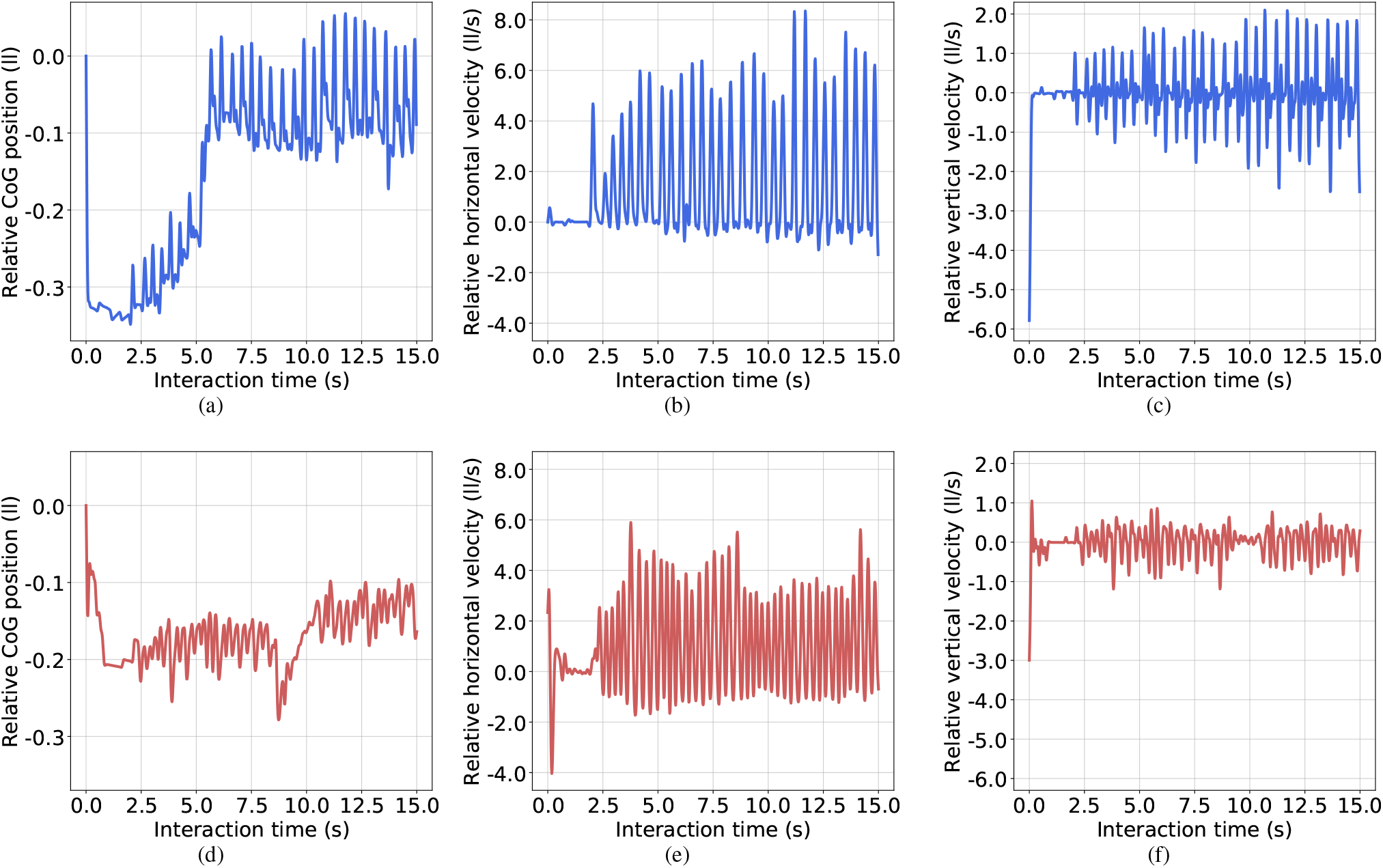
Gait analysis of best biped and best tripod. We pre-processed all data and divided by the leg length to eliminate the biped creature’s inherent advantage with longer leg length than the tripod creature. About the unit, *ll* represents the leg length, and *ll/s* represents the leg length per second. The upper three blue lines (a) (b) (c) are the relative Center of Gravity (CoG) position, relative horizontal and vertical velocities of the best biped while the lower red lines (d) (e) (f) are for the best tripod. Here, the relative CoG position is calculated based on the biped and tripod’s original CoG from the simulation’s beginning.

**Fig. 7:**
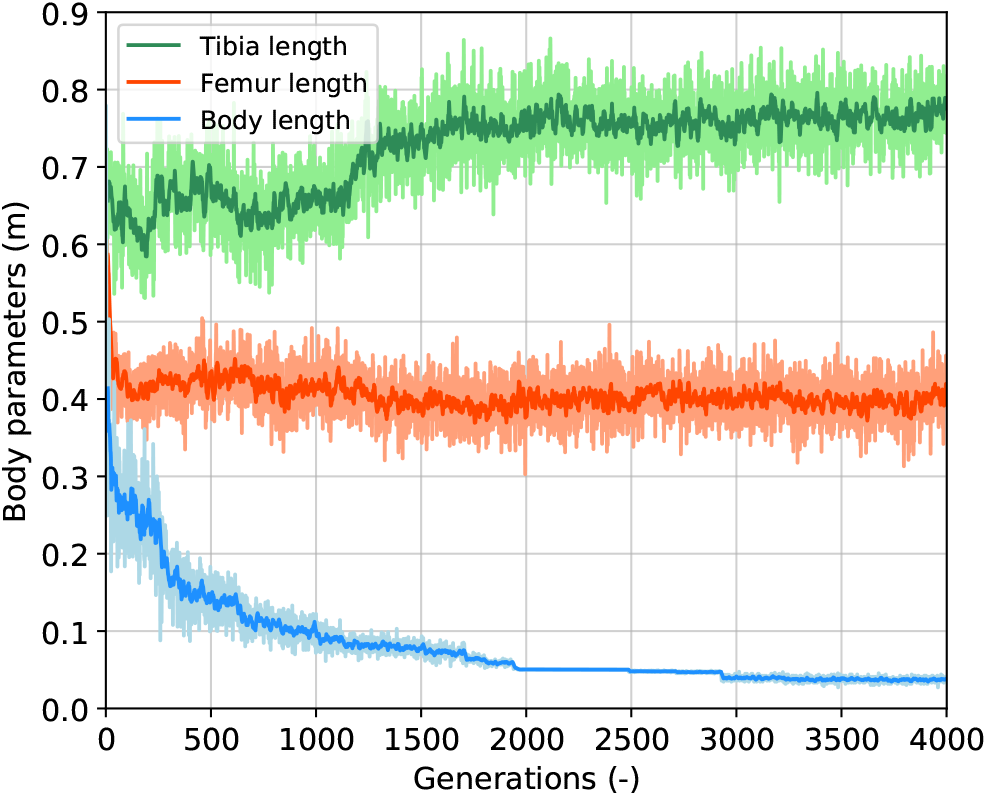
Parameters for the body length and the leg length over generations of the fast bipedal winner in the performancebased scaffolded case. Here, we use a smoothness function to draw the data clearly. The green curve and the orange curve are the tibia length and the femur length parameters, respectively. Their ranges are from 0.3 *m* to 0.9 *m*, and they are set to mutate freely. The light-blue curve is the body length parameter which is forced to decrease based on the current performance observed, and the range is from 0.05 *m* to 0.9 *m*.

The results described above were based on the best creatures, and those motivated us to create three cases (described at Section II.E) to understand the mechanisms leading to that difference. Initially, we wanted to identify the bipedal and tripodal creatures in our simulation, and we plot Fig. 8 to help us visualize the relationship between their gait and fitness. Over the course of 4000 generations for each run, these creatures present a strong tendency to evolve from tripod to biped while also drastically decreasing their body length. We run each case three times, and observe that tripod creatures rely on a bigger body to support their gait, never reaching the minimum length possible. After averaging the results from those three trials for each case, we plot Fig. 9, where we show the mean and standard deviation from each case. We observed that Case 1 (free scaffold) and Case 3 (performance-based scaffold) presented the bipedal results, while two-thirds of the runs from Case 2 (time-constrained scaffold) produced tripods as their best solutions.

**Fig. 8:**
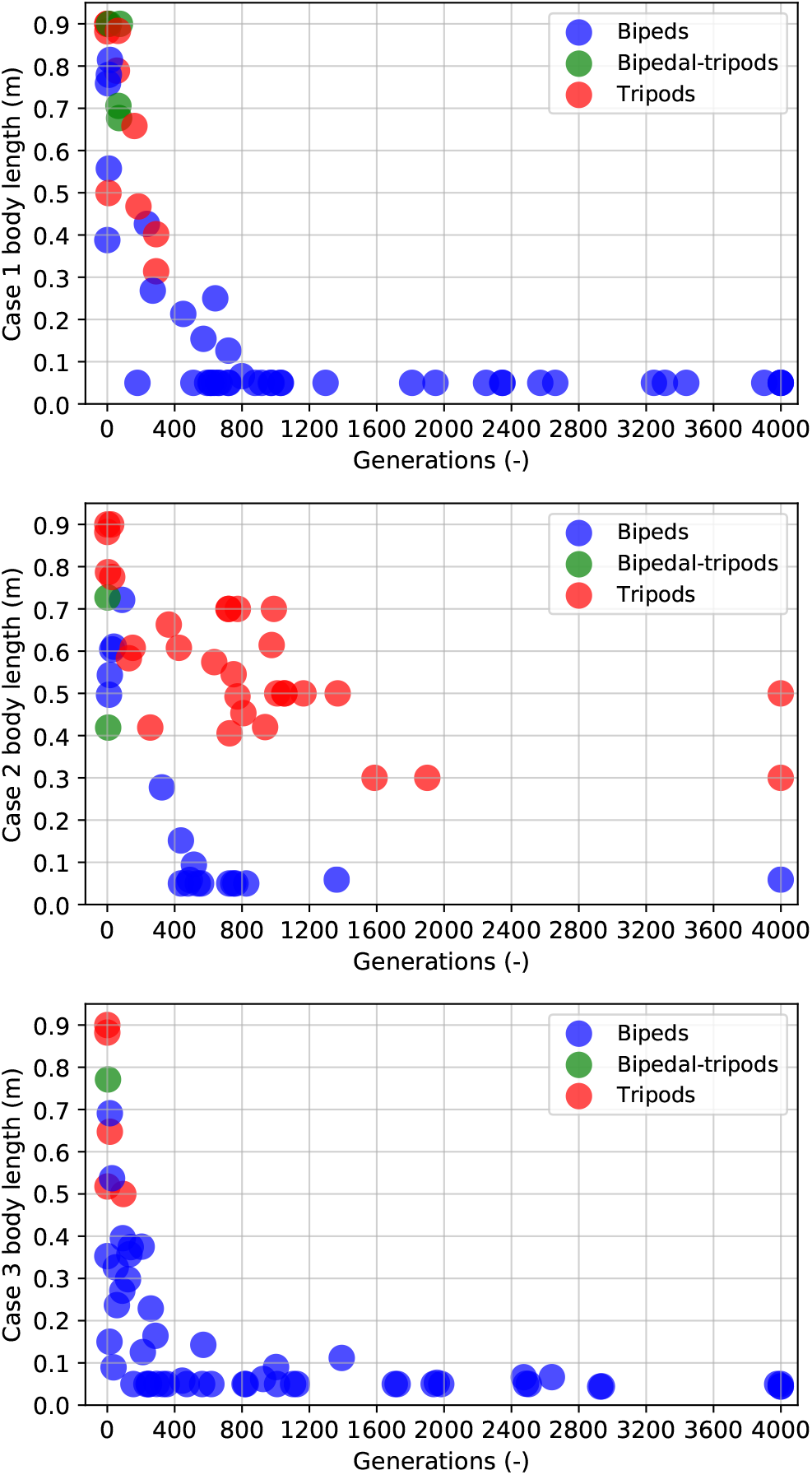
The scatter map of all three cases with different body lengths at every bump of the current fitness. a) Case 1: body length with free constraint during generations. b) Case 2: constraint body length with decreasing value during generations. c) Case 3: constraint body length based on the best fitness obtained so far during generations. Red stands for tripods, green for a hybrid bipedal-tripods, and blue for bipeds. Since the simulations started from a tripod individual with the longest body length, the scatter map is red in the beginning. By growing with different body length constraint mechanisms, most individuals become bipeds (blue) at the end.

**Fig. 9:**
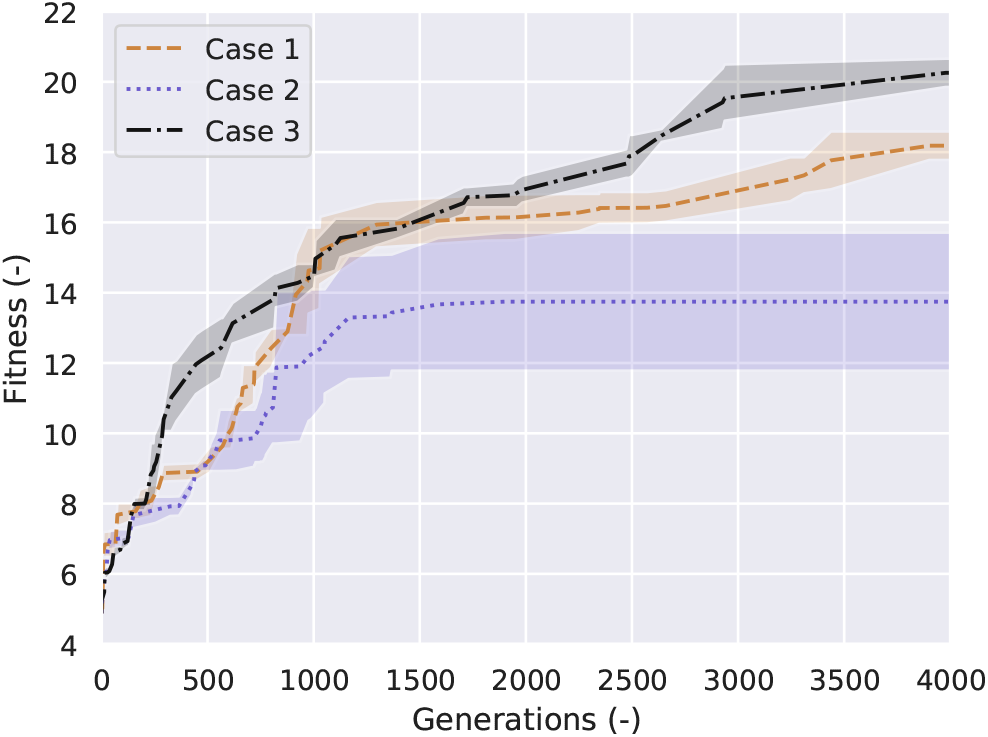
Results for all three cases within 4000 generations (three replicates for each case). The brown dashed, purple dotted, and black lines stand for Cases 1, 2 and 3, respectively.

In a comparison between Case 1 and Case 3 we can notice, from Fig. 9, that Case 3 has steeper progress in the first 800 generations, and ultimately reaches a higher fitness value (10% higher than Case 1). Case 2, on the other hand, struggles to evolve and only reaches 66% of the fitness from Case 1.

## IV. Discussion

### A. Body Length Supports Leg Growth

During the learning process of the best creature observed, the tripodal gait phase started with a long body and a short leg length, as shown in Fig. 7. The definition of long and short for the body length and the leg length was relative to the optimal body parameters. Upon analyzing our results, we found that the creature experienced two stages before achieving the optimal parametric combination of body and leg lengths. From 0 generations to 1000 generations, we observed a stability stage, with the body length rapidly decreasing from 0.9 *m* to 0.1 *m* while the tibia and femur lengths were near-constant. The second stage, marked by a gradual speed increase, is defined by an increase in the femur length while the body length reaches the minimum value set in the simulation.

In the first stage, the long body/short leg and its tripodal gait guaranteed the system to be stable to form a simple tripod control. Naturally, with an ever decreasing support, the short legs transition to a bipedal gait with a mature biped controller, and this triggers an increase in leg length to reach higher fitness values with an upright posture. This gait analysis allowed us to hypothesize on on the internal mechanism of a scaffolded learning approach, and also strongly agreed with the work from [7], where it is stated that roboticists could develop better systems by exploiting insights gained from studies on ontogenetic development. In this work, we state that A. stable tripodal gaits scaffold bipedal gaits and B. stable walking scaffolds speed increases, as seen in the transition from the first stage to the second, and our results are in strong agreement with a gait study with infants [10]. In this study toddlers who are still incapable of walking are supported on a treadmill and are capable of performing well-coordinated alternate stepping movements, in a very strong resemblance to an upright bipedal locomotion.

### B. Performance-based Scaffolds Bootstrap Learning

We proposed three cases of the scaffolded learning method in this paper. From the results shown in Fig. 8 and Fig. 9, we can state that the time-constrained scaffold (Case 2) hinders bipedal evolution, forcing the evolutionary process into a local optimum. On the other hand, a performance-based scaffold (Case 3) not only evolved bipedalism from tripods but also bootstraps its self-growth. Here, we set Case 1 as the baseline of independent bipedal walking learning. Although Case 1 is better than Case 2, the learning curve seen in Case 3 is the steepest. Case 3 takes the correct cues to transition from tripodal to bipedal, by shortening its body length gradually to allow the controllers to mature, in strong agreement to the results shown in [18], where the role of morphology in the control development is studied. Case 1 is free scaffolded learning, and the freedom provided prevents the system from assigning a higher priority on the contribution from the body parameter. Case 2 shows the negative effects of a poorly conceived support system, where the controller for the creature over-matures at a longer body length and stagnating at a tripod gait in most of the times.

We can analogize these cases to the human ontogenetic development for the performance of a cognitive task as a child. One is that parents deliberately do not interfere with a child’s learning, as seen in Case 1, another is that parents slowly reduce their assistance for this child based on their age, as seen in Case 2, and the other is that as this child performs this task parents adjust their support based on their perceived performance, as seen on Case 3. Broadening to pedagogical applications, Al Mamu et al. [6] provides a positive example of how to implement inquiry-based learning in an online environment, considering the lack of direct teacher or peer support. However, they mentioned that recent research rises more attention as challenges increase when adopting a free scaffold in the self-regulated learning environment without direct support from teachers. Therefore, only by choosing a suitable method can we effectively accelerate the learning process, which is in agreement with our work of physical robots [17], where we show the negative effects of an improperly enforced developmental process on a robot.

## V. Conclusions

In this paper, we introduced a scaffolded learning method on a creature capable of adapting its body and controller, hence bootstrapping a bipedal controller from a stable tripodal gait. Our results show that scaffolded learning with the right parameters is more productive than leaving a system free to learn independently. However, we also show that this is only true when the appropriate incentives behind scaffolded learning exist, effectively shortening the learning process with a performance-based scaffold, while a time-constrained scaffold is worse than the free learning case.

As the field of Robotics suffers from the curse of dimensionality and the Reality Gap, our proposed method should be used on robots for faster deployment of learning algorithms and a bottom-up construction of this knowledge base. The same concept explained herein could be transposed to a simulation-scaffolded reality, with the eventual removal of the training wheels to reproduce a reality-compatible behavior. As is the case with humans, robots should also be capable of using their previously acquired knowledge to aid their learning of complex tasks. After all, if Newton could see further, it was by standing on the shoulder of giants.

